# Increased susceptibility for pathogenesis of post-traumatic epilepsy in offspring exposed to deltamethrin during gestation and ameliorative effects of dietary curcumin

**DOI:** 10.1101/2023.12.03.569796

**Authors:** Prince Kumar, Kamendra Kumar, Deepak Sharma

## Abstract

Deltamethrin (DLT) is a most potent and widely used pesticide that does not cross Blood Brain Barrier (BBB) in adults. While it considered as safe, its lipophilic properties makes it a neurotoxic substance specially in early stages of brain development. It has shown neurotoxic effects on the brain by hyper-excitation of neurons. Epilepsy is a neurological disorder with recurring seizures where epileptogenesis occurs due to hyperexcitation of neurons. In various kinds of epilepsy, post-traumatic epilepsy (PTE) is a common epilepsy in children due to traumatic brain injury (TBI). PTE, however reportedly alleviated by curcumin in rats. Therefore, in the current study, we assessed the effect of gestational DLT exposure on the severity of PTE. The pregnant rats were injected with 0.75mg/kg-b/w of DLT dissolved in 1% DMSO each day of gestation between days 7-15. Epilepsy was induced four months postnatally, and curcumin was orally administered by oral gavage. ECoG, behavioral tests, Golgi Staining, immunofluorescence, and immunohistochemistry was performed to assess the pathogenesis, severity of epilepsy, and mitigating effects of curcumin. The results indicated the neurotoxic effects of DLT by raising the severity of seizures in an electrophysiological and behavioral manner. PTE decreased the dendritic branching and arborization. Sodium channel overexpression is an important reason for the hyperexcitation of neurons during the pathogenesis of epilepsy. DLT enhanced the increase in expression of both sodium channel subunits NaV1.1 and NaV1.6 during epileptogenesis. Similarly, synaptic markers PSD95 and SYP decreased. Astrocytic and microglial activation increased during pathogenesis of PTE. The antiepileptic effects of curcumin alleviated the effects on electrobehavioral response, neuronal arborization, and levels of NaVs, PSD95, SYP, GFAP and Iba1 in epilepsy. However, DLT raised the severity and susceptibility of epilepsy and decreased the antiepileptic effects on gestationally DLT-exposed epileptic animals. Our result demonstrates the gestational neurotoxic exposure of DLT increased the severity and susceptibility for PTE while decreasing the antiepileptic effects of curcumin.

## 1. Introduction

Deltamethrin (DLT) is a type II pyrethroid pesticide used in agriculture, aquaculture and household insecticides. Although DLT is typically considered a low-toxicity substance, excessive use can pose a potential risk to humanity. Due to its lipophilic nature, DLT can enter the mammalian system through different routes, including gastrointestinal, skin, and respiratory passages [1]. Notably, the known direct exposure is inhalation of fumes, and food/water contamination of DLT causes intoxication. An increasing number of studies show the accumulation of DLT in tissue of various organs in the body [2]. DLT is also reported to impair function in multiple organs, including numerous pathological changes and genetic defects [3]. Although DLT does not cross the BBB, DLT has still been reported to cause neurotoxicity in zebrafish [4]. DLT exposure’s effects are not evident in adults, yet its role in neurodevelopmental disorders is significant [5,6].

Epilepsy is a neurological disorder characterized by recurrent and unpredictable seizures. Epilepsy can occur due to various causes, such as genetics, brain tumors, stroke, and infection. However, occurrence after a head injury is a leading cause of epilepsy in children [7]. Epilepsy after a traumatic brain injury (TBI) is called post-traumatic epilepsy (PTE). Spontaneous reoccurring seizures are the most distinguished characteristic of PTE. Notably, the major cause of epileptogenesis from TBI is hyperexcitation of neurons due to changes in activity and expression channel proteins in neurons that subsequently causes neuronal death [7,8]. Consequently, causing behavioral alterations as well [9–11].

Curcumin is the main bioactive component of turmeric with potent anti-oxidant, anti-carcinogenic properties and neuroprotective activity. It is used as a traditional medicine to treat several clinical conditions like, inflammation, cancer, epilepsy and depression [12,13]. Growing evidence of literature has documented their neuroprotective efficacy in oxidative stress-induced brain insult [14–16], amyloid-beta-induced oxidative brain injury and 6-hydroxydopamine hemiparkinsonian model [17,18]. In epilepsy, multiple reports show its antiepileptic effects [19]. Increasing reports also show its role in alleviating the PTE effects after TBI in various PTE models [20–22]. These reports advocated the possible pleiotropic pharmacological potency of curcumin to treat complex disorders.

Various epilepsy models are employed in research, such as chemical models utilizing kainic acid and pilocarpine to imitate temporal lobe epilepsy (TLE) in rodents [23]. Other alternatives include the electrical stimulation-induced kindling model, genetic models involving voltage gate-linked mutations, and developmental models [24]. The controlled cortical injury model is utilized to replicate clinical scenarios in PTE. Still, it does not yet incorporate controlled focal seizures, which are necessary for comprehending pathogenesis and inflammation in epilepsy [25]. Additionally, the FeCl_3_-induced model connects epileptiform activity with vascular injury, which may encompass inflammation, oxidative stress, and vascular dysfunction. Since DLT is a potent source of neurotoxicity during brain development and the prevalence of PTE is high in children. It becomes an interesting question whether gestational exposure to DLT affects the susceptibility of epilepsy to developmentally exposed individuals. Therefore, the current study focuses on DLT exposure in pregnant female rats and later in their adult progeny, assessing where the DLT exposure during the embryonic stage affected the severity of PTE. The second aim of the current investigation is to address the alleviating effect of curcumin in gestationally DLT-exposed PTE-induced individuals.

## 2. Material and Methods

### 2.1. Test Species and Husbandry

Wistar rats used in the present study were bred and maintained in the Central Laboratory Animal Resources, Jawaharlal Nehru University. Care and maintenance of animals were carried out as per the guidelines laid by the Committee for the Purpose of Control and Supervision of Experiments on Animals (CPCSEA) and experimental protocols were approved by the Institutional Animal Ethics Committee (IAEC) of Jawaharlal Nehru University, New Delhi. The animal room temperature was maintained on standard conditions, i.e., 25±2°C, relative humidity of approximately 60 ± 10% and 12h light-12h dark photoperiod. Food and water were provided *ad libitum* throughout the study. The health status of each rat was checked by observing various criteria such as tail sores, posture hunch, grooming, scratching, rearing, red nose rim, red eye rims and tumors.

### 2.2. Breeding and treatment

For the breeding process, standard plastic cages (52 cm x 28 cm x 22 cm) were used to house two female rats and one male rat. The estrous cycle of the females was monitored each morning by collecting vaginal lavage on a glass slide using a Pasteur pipette filled with phosphate-buffered saline (PBS). The lavage samples were then observed under a microscope. Mating was confirmed by the presence of spermatozoa in the vaginal smear, and the day of confirmation was marked as Gestation Day zero (GD0). To the pregnant rats, a solution of DLT dissolved in 1% dimethylsulfoxide (DMSO) was injected intraperitoneally (i.p.) at a dosage of 0.75 mg/kg body weight per day during GD 7-15 (n=6). The animals were closely monitored for any signs of toxicity or changes in body weight as potential reactions to the treatment throughout the entire period until delivery. The day the pups were born was designated as postnatal day zero (PND 0). As a control group, an equivalent quantity of 1% DMSO was injected into control females (n=6), and the pups born to these females were used as age-matched controls. Only male rats were used in the present study to minimize sex-related issues. In adulthood, these animals were injected with FeCl_3_ stereotaxically to generate iron-induced epilepsy [11]. The curcumin (100 mg/kg bw/day in corn oil; Sigma) was administered orally (p.o.) for 30 days from the day of iron treatment. On the bases of iron and curcumin treatments, animals were further grouped as follows-1) DLT+Epileptic (DLT+Epi), 2) DLT+Epileptic+Curcumin (DLT+Epi+Cur), 3) Epileptic (Epi), 4) Epileptic+Curcumin (Epi+Cur) and 5) vehicle control (saline). Control, DLT+Epileptic, and Epileptic group animals also received corn oil as a vehicle (Fig.1).

**Fig. 1:**
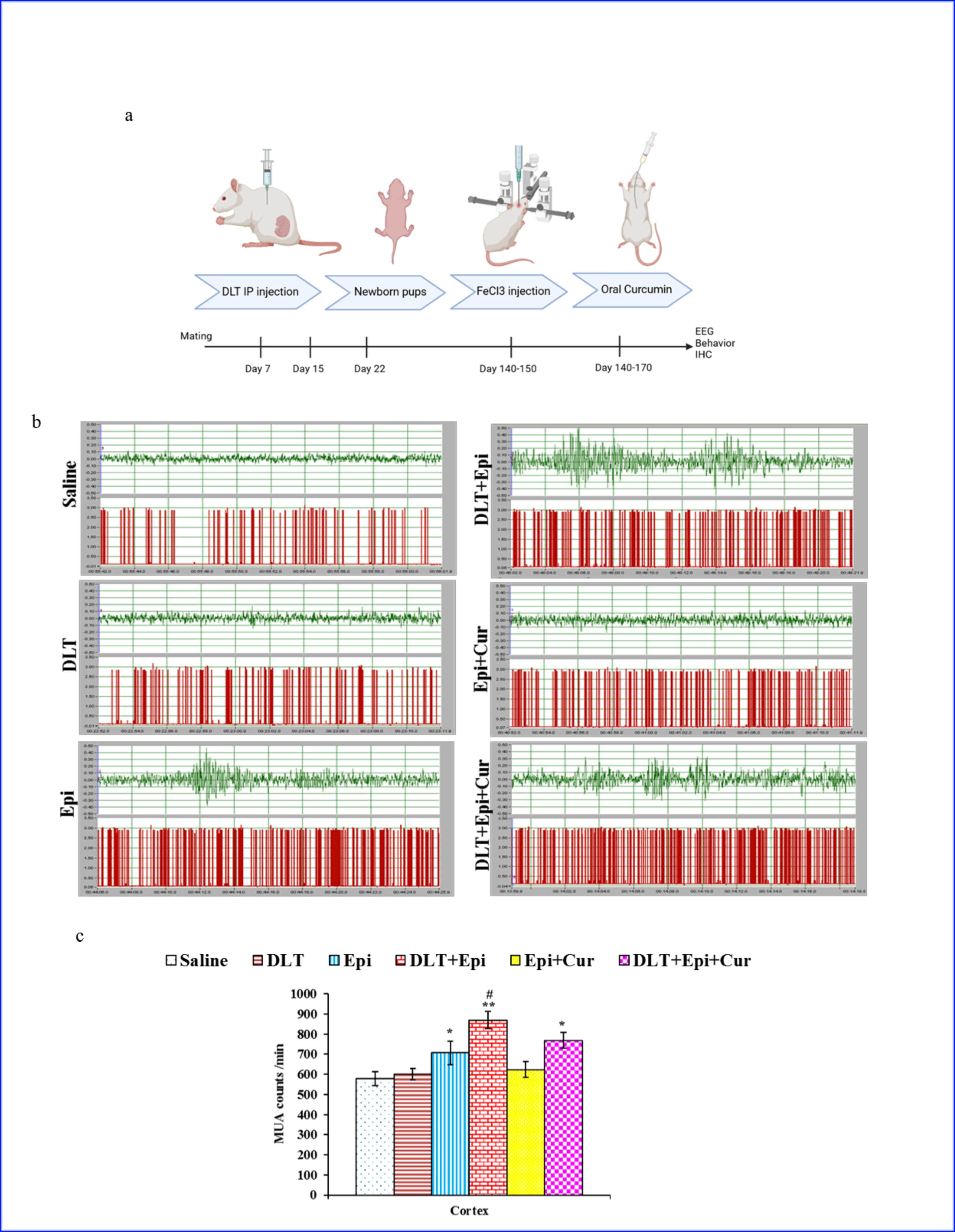
Strategy of the study to generate for DLT toxicity and inducing epilepsy treatment plan (a); further electrophysiological recordings showing patterns of ECoG and MUA count (b) of saline, DLT, Epi, DLT+Epi, Epi+Cur and DLT+Epi+Cur treated group animals. All the epileptic group presented an increase in the spike-wave complexes along with an increase in the MUA counts as compared to the saline group animals. The curcumin treatment prevented the epileptic firings and significantly decreased MUA counts. However, the curcumin treatment did not cause recovery from the increased epileptic firing and MUA counts in the DLT+Epi+Cur-treated group animals (c). No obvious difference was recorded in the ECoG and MUA count of saline and Epi+Cur-treated group animals. Values are presented as the mean ± SEM (n = 6).

### 2.3. Stereotaxic surgical procedure

Surgical procedures were performed as per the previously described method [21,26]. Under surgical anesthesia (4% isoflurane), animals were placed in a stereotaxic apparatus. A midline incision was made, the exposed skull was thoroughly cleaned, and burr holes (0.5 mm in diameter) were drilled for iron injection and electrode placement. Solution of FeCl_3_ (5 µl of 100 mM)/saline (pH adjusted) was injected at a fixed rate 1µl/min using Motorized Stereotaxic Nano-injector (Stoelting, USA) in the somatosensory region (coordinates: 1.0 mm anteroposterior, 1.0 mm lateral, and 2.0 mm ventral from bregma) through a burr hole. After injection, the burr hole was sealed with bone wax to prevent the outflow. Further, four epidural screw electrodes were placed, 2 mm posterior and anterior to bregma and 2 mm lateral to the midline, surrounding the somatosensory region for the electrocorticogram (ECoG) and Multi-unit activity (MUA) recordings. In addition, one screw electrode was also placed on the midline over the frontal sinus to serve as a reference. The free ends of all seven electrodes soldered successively to a 9-pin adapter and the entire assembly was fixed tightly on the skull using dental acrylic. The electrodes and wires used in the present study were tissue compatible and procured from P1 Technologies, VA, USA. Operated rats were provided with optimal post-operative care throughout the experiments. All the experimental animals were habituated in the recording chamber for three days prior to final electrophysiological recordings (ECoG and MUA).

### 2.4. In-vivo recordings

Electrocorticography and MUA were recorded as described previously [21]. MUA recording can provide a relatively large amount of data with a high spatial and temporal resolution. Along with ECoG recording, MUA can verify hyperexcitation and increased neuronal firing, ensuring the actual epileptic activity rather than arbitrarily recorded potential. Consequently, neuronal hyperexcitation firing causes synchronously raised MUA bursts laterally with ECoG paroxysms. Thus MUA counts were used for quantifying epileptic seizure activity [27]. The data was acquired using polyVIEW 16 (version 1.1) data acquisition system (Grass Technologies, USA) with model P511 high-performance AC pre-amplifiers (MODEL L AND LATER). Complex ECOG signals were amplified, filtered (300 Hz to 10 kHz) by high-performance AC pre-amplifiers, and electronically discriminated using a window discriminator (WPI, USA) to generate MUA recordings. The recordings were restricted to the awake static state [8].

### 2.5. Behavioral Tests

#### 2.5.1. Open Field Maze

An open-field maze is used for quantifying locomotory and animal anxiety-like behavior. The test is based on the wall-enclosed premise of a square shape (70cm x 70cm x 106cm) of an unknown environment for animals from which escape is prevented. Rodents spontaneously prefer the periphery of the apparatus to activity in the central parts of the open field. The test was performed as previously described [11]. In the current study, the maze consisted of 12 peripheral and four central squares. Rats were primarily placed in the center of the maze to explore it. The maze field was cleaned thoroughly to remove any previous animal scent clue using 0.1% acetic acid solution.

The parameters evaluated were locomotor or ambulatory activity by total distance traveled and mobile time. Vertical locomotor activity or rearing frequency is defined as the number of times the animals stand on their hind legs. The test was performed for five consecutive days with six trials of 3min each per day per animal. ANY-maze behavioral-tracking-software (Version 5.1; Stoelting, USA) was used for tracking analysis of video recording of the test.

#### 2.5.2. Three Chamber test

The Three-Chamber test is used for assessing cognition behavior in form of sociability, exploration of social novelty, and social gender preferences. The difference in this behavior in various CNS disorders makes it a pivotal test. In the present study, the test was performed as discussed previously [28,29]. The apparatus consisted of identical chambers (31.5 x 31.5 x 31.5 cm^3^) of transparent plexiglass connected by 10×10 cm^2^ doors. The rats were habituated 10 min for four days by housing animals in the empty cage and with age-matched conspecific animals of both genders for familiarization.

For sociability test, after the habituation period, the rats were guided to the central chamber of the apparatus, and the openings were closed to restrict their movement. In one of the side chambers, an unfamiliar rat was placed inside a bar-wired corral, while the other side chamber remained empty. Subsequently, the doors separating the chambers were opened, initiating the sociability test. Each experimental session had a duration of 10 minutes and was meticulously recorded using an overhead video camera.

For Social preference test, rats were led to the middle chamber and the openings in the apparatus were closed. A familiar female rat was then placed in a bar-wired corral in one of the side chambers, while a familiar male rat was placed in corral of the other side chamber. Similarly, an unfamiliar rat of same sex was placed in corral of one side of the chamber while the familiar rat was placed in corral of another side. While the test rat was placed in central chamber.

### 2.6. Tissue collection and processing

After electrophysiological recordings and behavioral analysis, all the animals were perfusion-fixed (transcardially) with 4 % paraformaldehyde prepared in 0.01 M phosphate buffer (pH 7.4). Post-fixed in the same fixative, cryo-protected in sucrose gradients, i.e., 10, 20 and 30 %, coronal sections (15 µm) were sliced using a Leica cryotome (CM1860 UV), collected on gelatin-coated slides and stored at -20 °C.

### 2.7. Golgi Staining

The Quick Golgi-cox staining procedure was conducted using a previously described method [30]. Following perfusion with either PBS alone or PBS followed by rapid-Golgi fixative solution (P-RGF), the brains were extracted and the cerebral cortex and cerebellum were removed. From both P-PBS and P-RGF groups of rats, tissue blocks of approximately 1 cm3 were post-fixed in RGF and impregnated with a 0.75% aqueous AgNO3 solution using various combinations (n=3) of post-fixation durations of 12, 24, 36, or 48 hours, followed by AgNO3 impregnation durations of 24, 36, or 48 hours. After the respective durations in AgNO3 solution, the tissues were washed thoroughly in 70% alcohol and 100 μm thick sections were prepared using a Leica cryotome (CM1860 UV). The prepared slides were visualized using a Leica DM6000 microscope equipped with a Leica DFC 310 FX digital camera and the LAS V4.2 multi-focus module. The module captured a series of images at different focus positions across the depth of the neurons, which were then combined into a single image containing all the in-focus neuronal processes in the stack. The spine density of a total of 220 neurons (110 Purkinje and 110 pyramidal neurons) was analyzed.

### 2.8. Immunofluorescence

Cryotome sections through cortical region were selected and processed for the double-immunofluorescence labeling with anti-synaptophysin (SYP), anti-PSD95 and anti-NF200. Sections were washed in phosphate-buffered saline (PBS). Followed by permeabilization with 0.5% Triton X-100 and non-specific protein blocking with 5% normal serum (dissolved in washing buffer), sections were incubated with the cocktail of primary antibodies, anti-synaptophysin (1:500, rabbit polyclonal, Santa Cruz) and anti-NF200 (1:1000, mouse monoclonal, Abcam) or anti-PSD95 (1:500, rabbit polyclonal, Invitrogen), anti-Nav1.1 (1:100, Alomone, Labs, Jerusalem, Israel; rabbit polyclonal), anti-Nav1.6 (1:100, Alomone Labs, Jerusalem, Israel; rabbit polyclonal) for overnight at 4 °C in a humid chamber. Next day, the sections were washed with PBS, and incubated with the cocktail of fluorochrome-conjugated secondary antibodies, anti-rabbit TRITC tagged (1:200, Sigma) and anti-mouse FITC tagged (1:200, Sigma) at room temperature in dark for 90 min. The sections were washed in four changes of PBS and mounted with VECTASHIELD® HardSet™ mounting medium with DAPI from Vector. The sections were visualized under a fluorescence microscope (Nickon Eclipse 90i) fitted with a digital camera DS-Qi1Mc. The images were acquired using NIS Elements AR version 4.0 software.

### 2.9. Immunohistochemistry

Immunohistochemistry using cryosections was performed to identify GFAP and Iba1 positive glial cells, as previously described in literature (cite). Briefly, the stored sections were taken out and left at room temperature (37 °C) for 1 hour before denaturation in 1% Triton-100. The sections were then incubated in the dark with 1% hydrogen peroxide to inhibit any endogenous peroxidase, followed by incubation in 10% normal goat serum (NGS) at room temperature for 90 minutes. Primary antibodies against anti-GFAP [1:500, sigma] and anti-Iba1 [1:1000; Abcam] were added and the sections were incubated overnight at 4 °C in a humid chamber. The next day, the slides were incubated at room temperature for 1 hour and then washed three times with PBS. The sections were then incubated with HRP-labeled secondary antibody (IgG 1: 100) at room temperature for 2 hours and diaminobenzene (0.25% in 1% H2O2) dissolved in PBS was added for 10-15 minutes. Finally, the sections were observed under an Olympus microscope, and three sections per rat were used for quantification. The expression of GFAP and Iba1 was calculated using the ImageJ software by measuring the area under the curve.

### 2.10. Statistical analysis

All results are expressed as mean ± standard error of means (SEM) and were analyzed by one-way analysis of variance (ANOVA). Tukey’s test was used as the post hoc comparison. The significant level was set at p<0.05. Sigma Stat 3.5 software was used for statistical analysis.

## 3. Results

### 3.1. In-vivo Electrophysiology

In order to determine the influence of DLT exposure on the development of epileptic seizures, according to study plan (Fig. 1a), rats were subjected to the ECoG and MUA recordings at 30-day post-FeCl_3_-injection. Epilepsy induced animals group developed abnormal spikes of ECoG characterized by polyspikes and paroxysms of spikes wave complexes of variable intervals (Fig. 1b). Simultaneously recorded MUA count also increased in numbers as abnormal spiking pattern was observed in epileptic group, thus confirming the epileptic field potential instead of randomly recorded potentials of nearby structures. Quantitative analysis of MUA counts depicted a significant increase of epileptiform electrographic activity and MUA count in all epileptic groups, i.e., Epileptic (*p* < 0.05), DLT+Epi (*p* < 0.01) and DLT+Epi+Cur (*p* < 0.05) groups as compared to the age-matched saline group animals. However, gestational DLT treatment caused a significant increase in MUA count compared to the Epileptic-only group (Fig. 1c). In addition, DLT+Epi+Cur group animals also presented a significantly increased in MUA count compared to the Epi+Cur treated group animals (*p* < 0.05). No significant difference in MUA count was observed between the saline and gestationally DLT-exposed saline-injected group. These results confirm that curcumin treatment attenuated the iron-induced epileptic discharge in the epileptic group animals but failed to restore epileptic firing in the gestationally DLT-exposed epileptic group animals.

### 3.2. Behavioral Tests

The alteration in the behaviors of the animals was assessed through OFT and Three chambered tests. In cases of epilepsy, the OFT experiment showed a significant increase in the distance traveled (Fig. 2a), mobile time (Fig. 2b), and rearing activity (Fig. 2f) of animals (*p* < 0.05). Conversely, there was a significant decrease in the immobile (Fig. 2c) and freezing time (Fig. 2e) of rats (*p* < 0.05). Similarly, in DLT-exposed epilepsy-induced animals, i.e., DLT+Epi, there was a significant increase in distance traveled, mobile time, and rearing activity. In contrast, a decrease in the freezing time of rats was observed in the maze (*p* < 0.05). Curcumin treatment in both Epileptic only and DLT-exposed epileptic, e.i., Epi+Cur group and DLT+Epi+Cur groups, showed decreased distance traveled, mobile time, and rearing as compared to individual comparable groups for Epileptic and DLT+Epi. Similarly, immobile time, stereotypic time, and freezing time increased when compared to their comparable groups (*p* < 0.05). Additionally, the path traveled by rats shows decreased thigmotaxis in DLT-exposed epileptic rats (Fig. 2g). However, alleviation in thigmotaxis was observed in curcumin-treated rats of both epileptic as well as DLT-exposed epileptic groups. The Three Chamber test depicted that epileptic group animals spent significantly more time either with the empty cage or with the unfamiliar rat than the saline-treated group animals, indicating a decreased social novelty (*p* < 0.05 & 0.01) (Fig. 3a, b) and social preference (*p* < 0.05)(Fig. 3c, d). The curcumin treatment counteracted the epilepsy-mediated impairments by restoring the preference for the novel rat (*p* < 0.05, 0.01). However, the curcumin treatment did not recover the preference in the DLT+Epi animals (*p* < 0.01, 0.01). Suggesting severity is more pronounced in the DLT+Epi group animals. Further, Epileptic, DLT+Epi, and DLT+Epi+Cur treated group animals presented a significantly reduced social novelty index (p < 0.01, 0.01) (Fig. 3e, f) and social preference index (*p* < 0.01, 0.01) (Fig. 3g, h) as compared to the saline and Epi+Cur treated group animals. Conversely, animals from the saline group and Epil+Cur treated group showed no significant difference in the social novelty index (Fig. 3e, f) and social preference index (Fig. 3g, h).

**Fig. 2:**
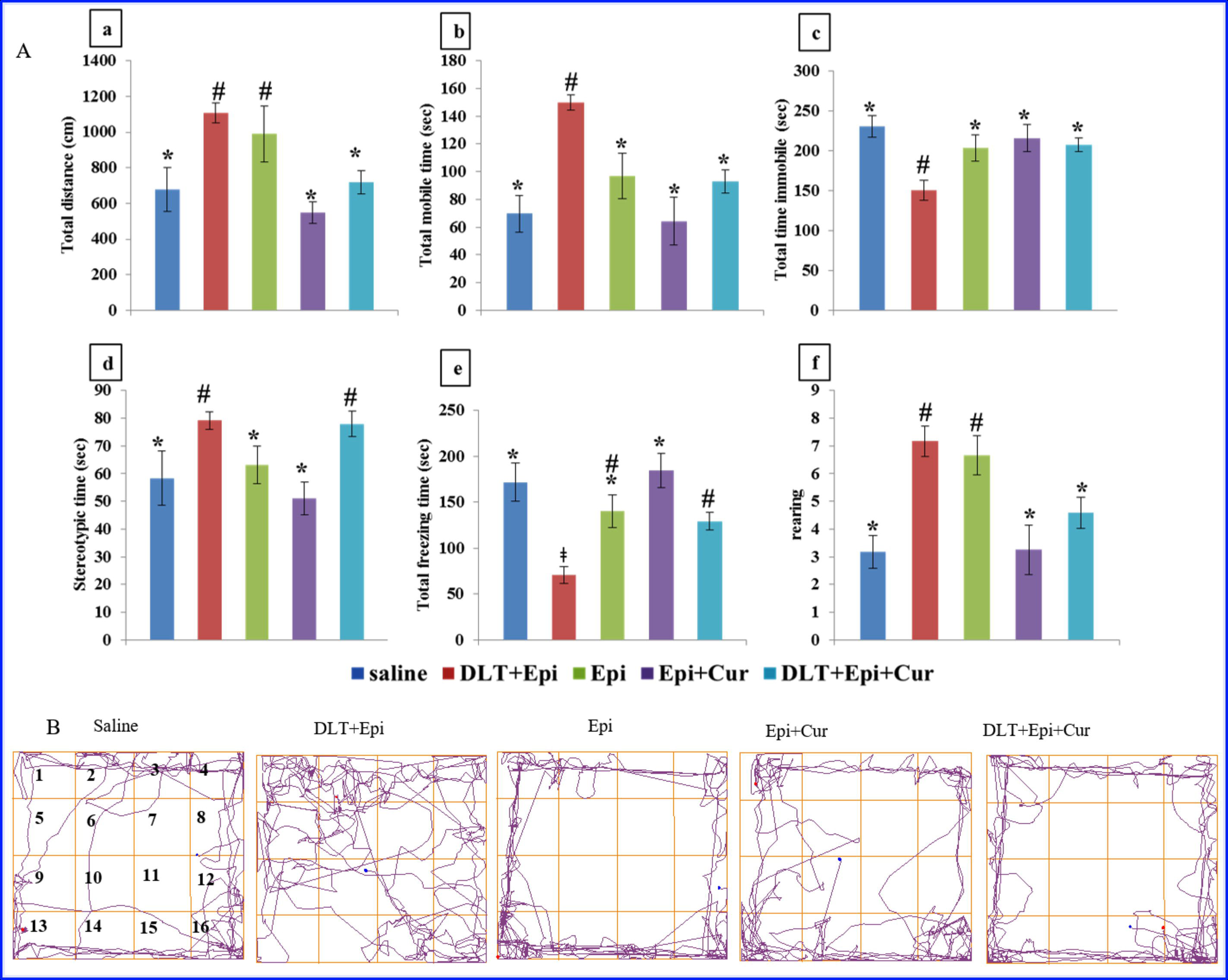
Bar graph demonstrating the total distance traveled (a), total time mobile (b), total time immobile (c), stereotypic time (d), total freezing time (e) and rearing (f) and pattern of traveled path by rats (g) in Open field test of saline, Epileptic, DLT+Epileptic, Epileptic+Curcumin and DLT+Epileptic+Curcumin treated group animals at 30-day post iron treatment. Values are presented as the mean ± SEM (n = 6). Values with the same symbol within each bar are not significantly different at the 5% level of probability.

**Fig. 3:**
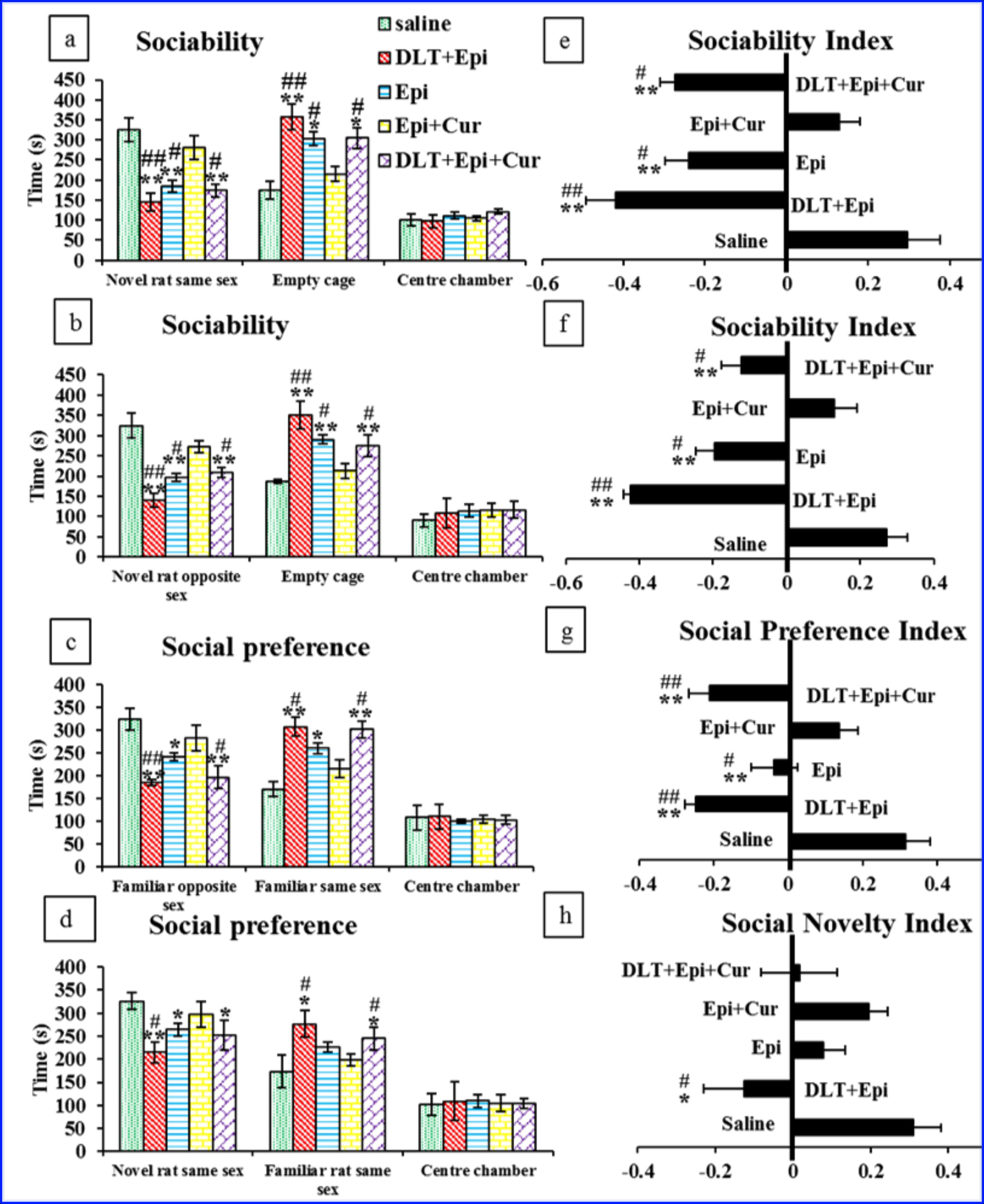
All the iron-induced epileptic group animals spent significantly more time either with the empty cage or with the unfamiliar rat than the saline-treated group animals, indicating decreased social novelty (a, b) and social preference (c, d). The curcumin treatment was able to counteract the epilepsy-mediated impairments by restoring the preference for the novel rat. However, the curcumin treatment did not recover the preference in the DLT+Epi group animals. Suggesting severity is more pronounced in the gestationally DLT-exposed iron-induced epileptic group animals. Further, Epileptic, DLT+Epileptic and DLT+Epi+Cur treated group animals presented a significantly reduced social novelty index (e, f) and social preference index (g, h) as compared to the saline and Epileptic +Curcumin treated group animals. Conversely, animals from saline group and Epileptic+Curcumin treated group showed no significant difference in the social novelty index (e, f) and social preference index (g, h). \*\**p* < 0.01, \**p* < 0.05, saline vs. Epi, DLT+Epi, DLT+Epi+Cur; ##*p* < 0.01, #*p* < 0.05, Epi+Cur vs. Epi, DLT+Epi, DLT+Epi+Cur (One-way ANOVA followed by post hoc Tukey’s test for multiple comparisons among groups).

### 3.3. Cortical dendritic complexity

Further dendritic degeneration due to epileptic seizures in cortical neurons is examined by Golgi staining. Golgi staining made individual neurons in layer III, layer V, and layer VI neurons visible in the saline, Epi, DLT+Epi, Epi+Cur, and DLT+Epi+Cur (Fig. 4a). Similarly decrease in the dendritic spine was spotted in Epi and DLT+Epi group (Fig. 4b) The altered morphology of dendritic tree in terms of reduced branching with short primary dendrites and terminal branching was recorded in the Epi and DLT+Epi group animals. Curcumin supplementation in epileptic animals prevents a decrease in dendritic branching. However, no effect of curcumin administration on reduced dendritic branching was observed in DLT+Epi+Cur. In addition to dendritic branching, similar results were reported in the spine density measurements, where the spine density was significantly decreased in the layer III, layer V, and layer VI of Epi (*p* < 0.05), DLT+Epi (*p* < 0.01) and DLT+Epi+Cur (*p* < 0.05) as compared to the saline-treated group animals (Fig. 4b and c).

**Fig. 4:**
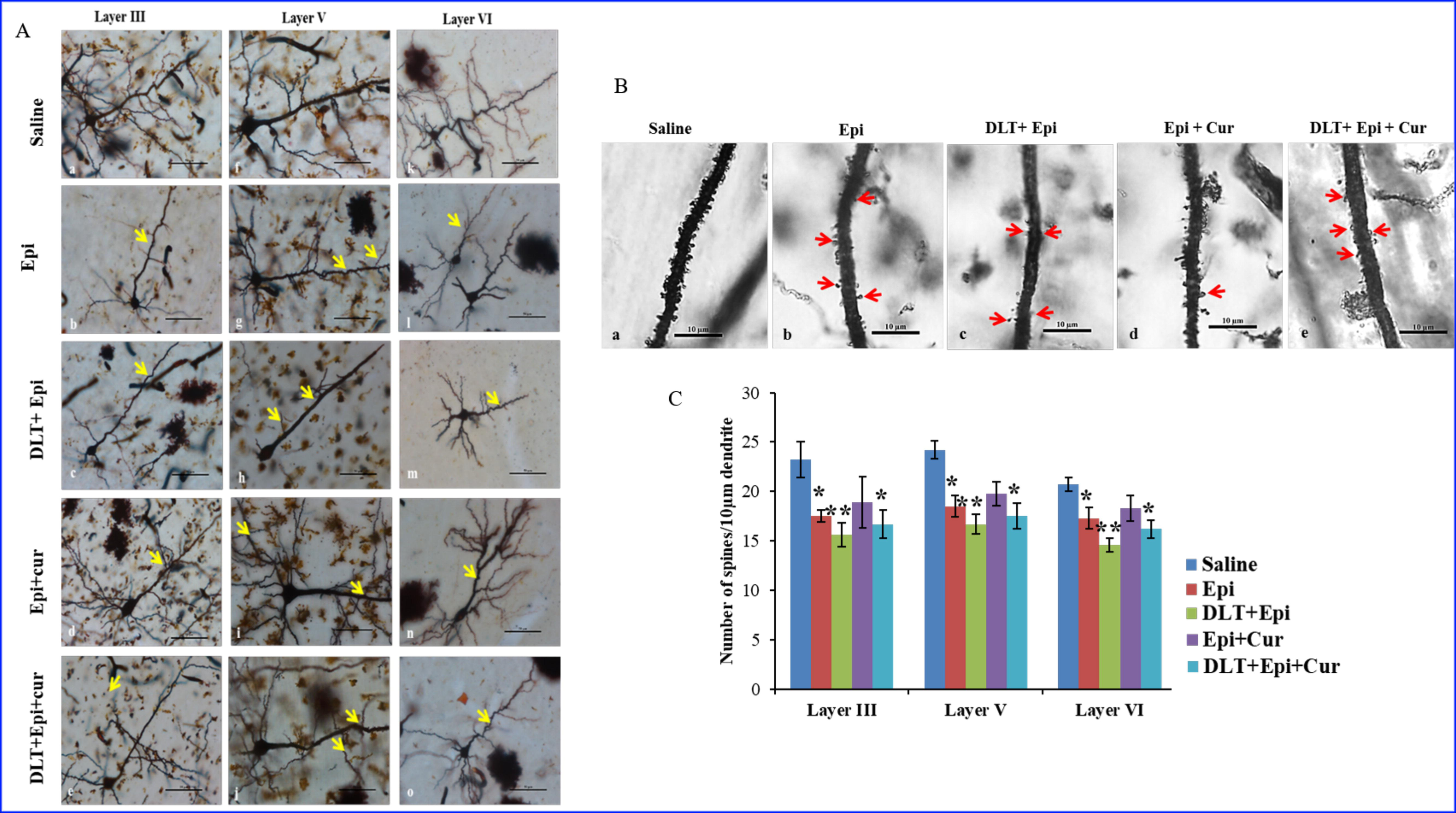
Representative images of Golgi stained neurons from the cortical layer-III, V and VI of Saline, Epileptic, DLT+Epi, Epi+Cur and DLT+Epi+Cur treated group animals. Arrows indicate the rapid-Golgi-stained pyramidal neurons with distorted morphology. A significant decrease in dendritic arborization in the epileptic group is demonstrated (a). The scale bar = 50μm. Graphs showing the spine density (mean ± SEM) measurement of the cortical neurons from all the groups. All the epileptic group animals presented a significant decrease in the spine density of cortical neurons as compared to the saline and Epi+Cur treated group animals (b). However, the severity is more pronounced in the gestationally DLT exposed epileptic group animals as the spine density decreased drastically. Values are presented as the mean ± SEM (n = 6).

### 3.4. Immunofluorescence detection of post and pre-synaptic neurons

Postsynaptic density protein (PSD95) is a marker of postsynaptic neurons that provides information about synapsis, while Neurofilament (NF) is a marker of neuronal axon. Here, we determined the localized expression of PSD95 in the Cortical neurons. We performed double immunostaining of anti-PSD95 with anti-NF in all of the aforesaid groups. Epileptic group animals presented a significant decline in the expression of postsynaptic density marker protein (*p* < 0.05) as compared to the saline group animals. The decline was more robust in the DLT-exposed epileptic group animals (*p* < 0.001). The curcumin treatment prevented the loss of PSD95 (*p* < 0.05) proteins in Epi+Cur group when compared to the Epileptic-only group. However, the protective effect of curcumin was not evident in the DLT-exposed epileptic i.e. DLT+Epi+Cur group animals (*p* > 0.05) as compared to DLT+Epi group. However, even after curcumin treatment significant difference in PSD95 was found when DLT+Epi+Cur group compared to the saline group (Fig. 5a and b).

**Fig. 5:**
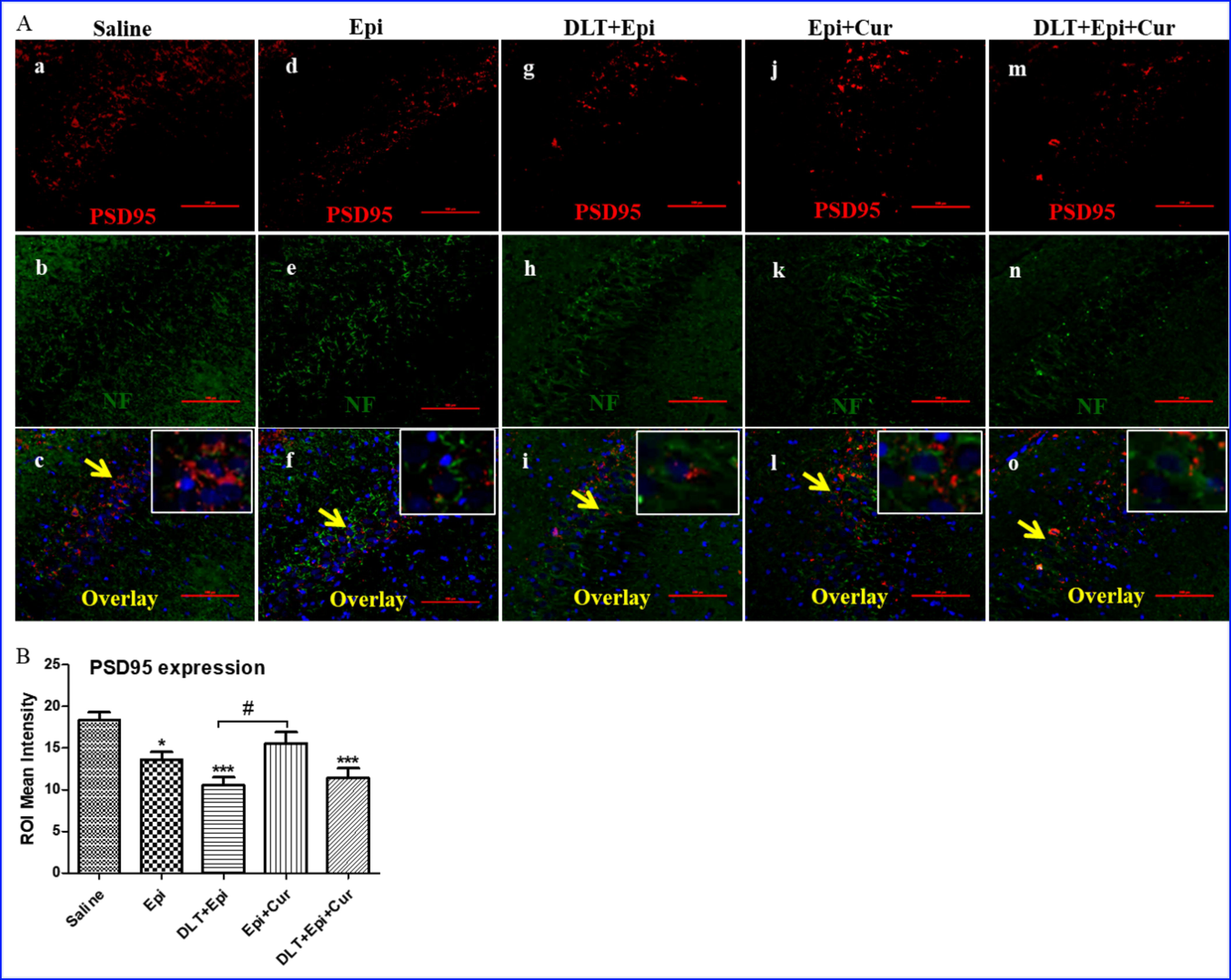
Double immunostaining of rabbit anti-PSD95 (detected with TRITC, red) and mouse anti-NF (detected with FITC, green) in the cortex of saline, Epi, DLT+Epi, Epi+Cur and DLT+Epi+Cur treated groups at 30-day post FeCl_3_ injection (a) and quantification of mean intensity of PSD95 (b). Insets showing the magnified images of double-labeled PSD95 and NF. Nuclei were stained with DAPI (blue). The scale bar = 100μm. Values are presented as the mean ± SEM (n = 6).

A similar trend was also reported in the expression pattern of presynaptic marker protein (SYP), where the expression of SYP was significantly low in Epi (*p* < 0.001) and DLT+Epi (*p* < 0.001) and DLT+Epi+Cur (*p* < 0.001) treated group animals as compared to the saline. Curcumin administration significantly prevented the loss of SYP in the iron-induced epileptic group animals (*p* < 0.05). In addition, SYP expression was also significantly low in Epi (*p* < 0.001), DLT+Epi (*p* < 0.001), and DLT+Epi+Cur (*p* < 0.001) group as compared to the Epi+Cur group. Shows that curcumin failed to alleviate the epilepsy-associated downregulation of SYP. However, no significant difference was reported in the expression of NF protein among the groups (Fig. 6a and b).

**Fig. 6:**
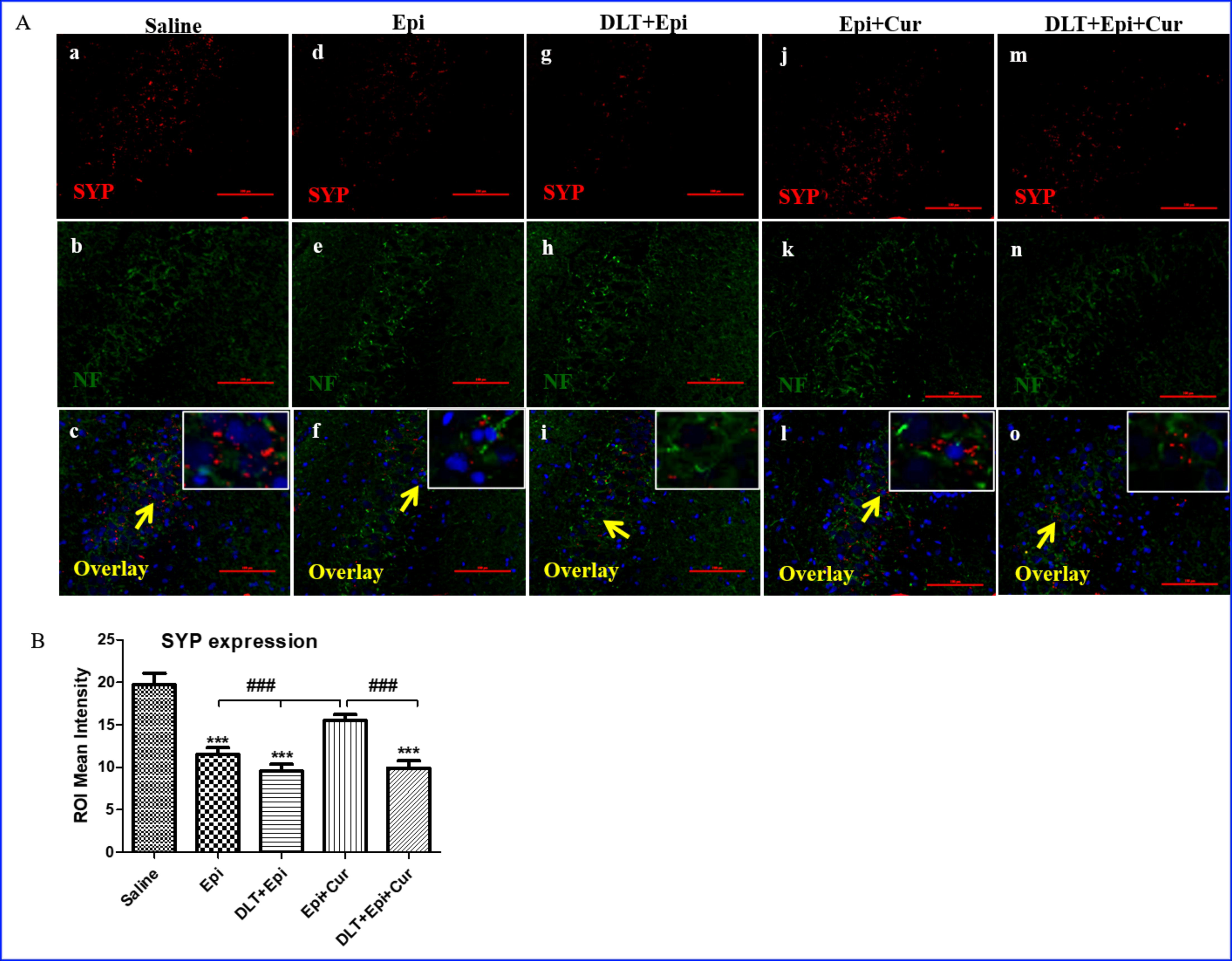
Double immunostaining of rabbit anti-SYP (detected with TRITC, red) and mouse anti-NF (detected with FITC, green) in the cortex of saline, Epileptic, DLT+Epi, Epi+Cur and DLT+Epi+Cur treated groups at 30-day post iron treatment (a) and quantification of mean intensity of SYP (b). Insets showing the magnified images of double immunostained SYP and NF proteins. Nuclei were stained with DAPI (blue). The scale bar = 100μm. Values are presented as the mean ± SEM (n = 6).

### 3.5. Immunofluorescence detection of Sodium channel subunits

In order to determine the regulation of sodium channel expression during epileptogenesis, the expression of Nav 1.1 and Nav 1.6 was quantified by double-labeled immunofluorescence staining. The findings indicate that both VGSCs (Nav1.1 and Nav1.6) were present in a co-expressed state. The results showed that epileptic group animals presented a significant rise in the expression of NaV1.6 and NaV1.1 (*p* < 0.05) compared to the saline group animals. The increased level was significantly higher among animals in the epileptic group exposed to DLT e.i. DLT+Epi (*p* < 0.05). By curcumin treatment, the overexpression of both sodium channel subunits is effectively prevented in epileptic group (Epi+Cur) animals as compared to the saline (*p* < 0.05). However, the results suggested that curcumin failed to produce any noticeable improvement in epileptic animals exposed to DLT, when compared to the saline group (Fig. 7a and b). Also, it was observed that both NaVs in Epi+DLT+Cur and Epi+Cur groups exhibited significant differences in the data (*p* < 0.001, *p* < 0.001).

**Fig. 7:**
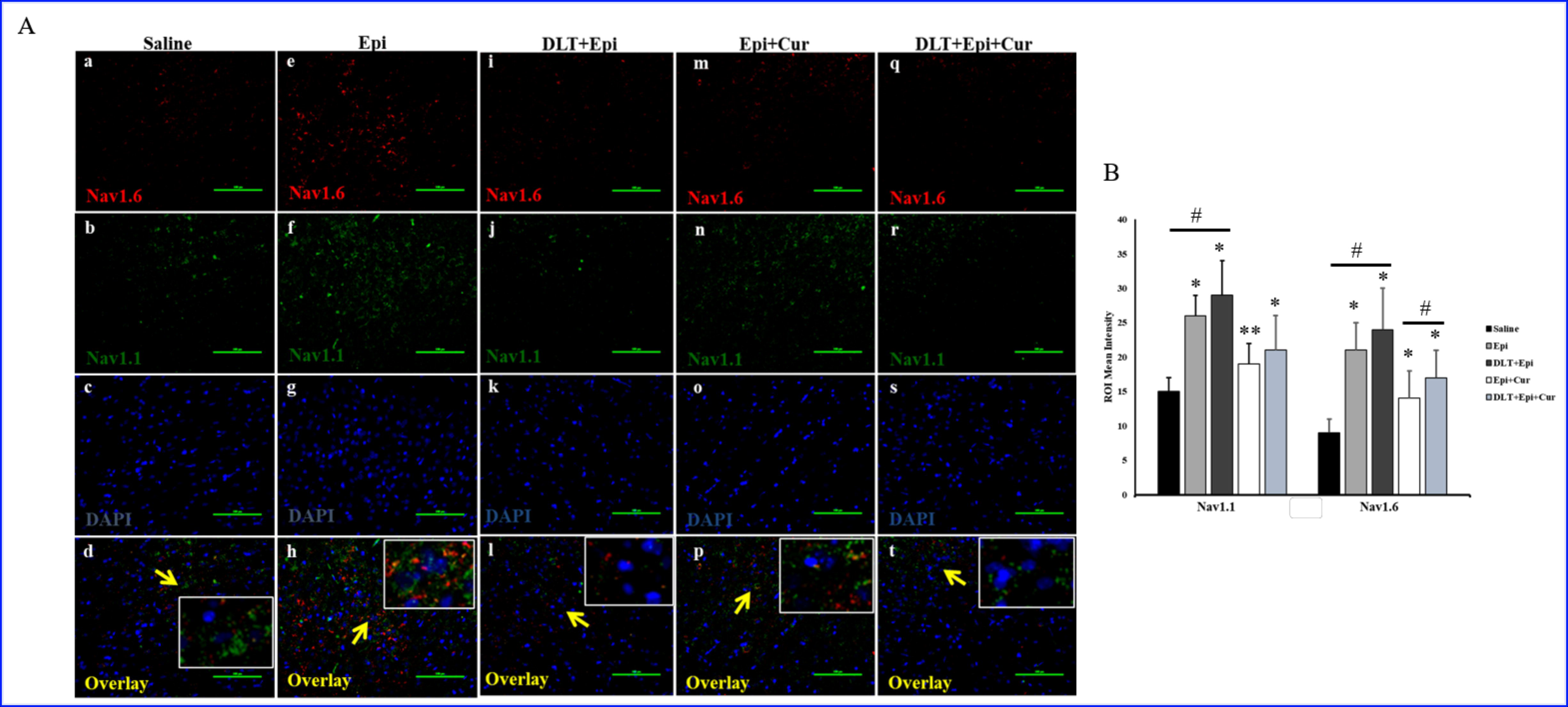
Double immunostaining of rabbit anti-Nav1.6 (detected with TRITC, red) and mouse anti-Nav1.1 (detected with FITC, green) in the cortex showing saline, Epi, DLT+Epi, Epi+Cur and DLT+Epi+Cur treated groups at 30-day post iron treatment (a). Quantification of mean intensity of Nav1.1 and Nav1.6 (b). Insets showing the magnified images of double-labeled Nav1.1 and Nav1.6. Nuclei were stained with DAPI (blue). The scale bar = 100μm. Values are presented as the mean ± SEM (n = 6).

**Fig. 8:**
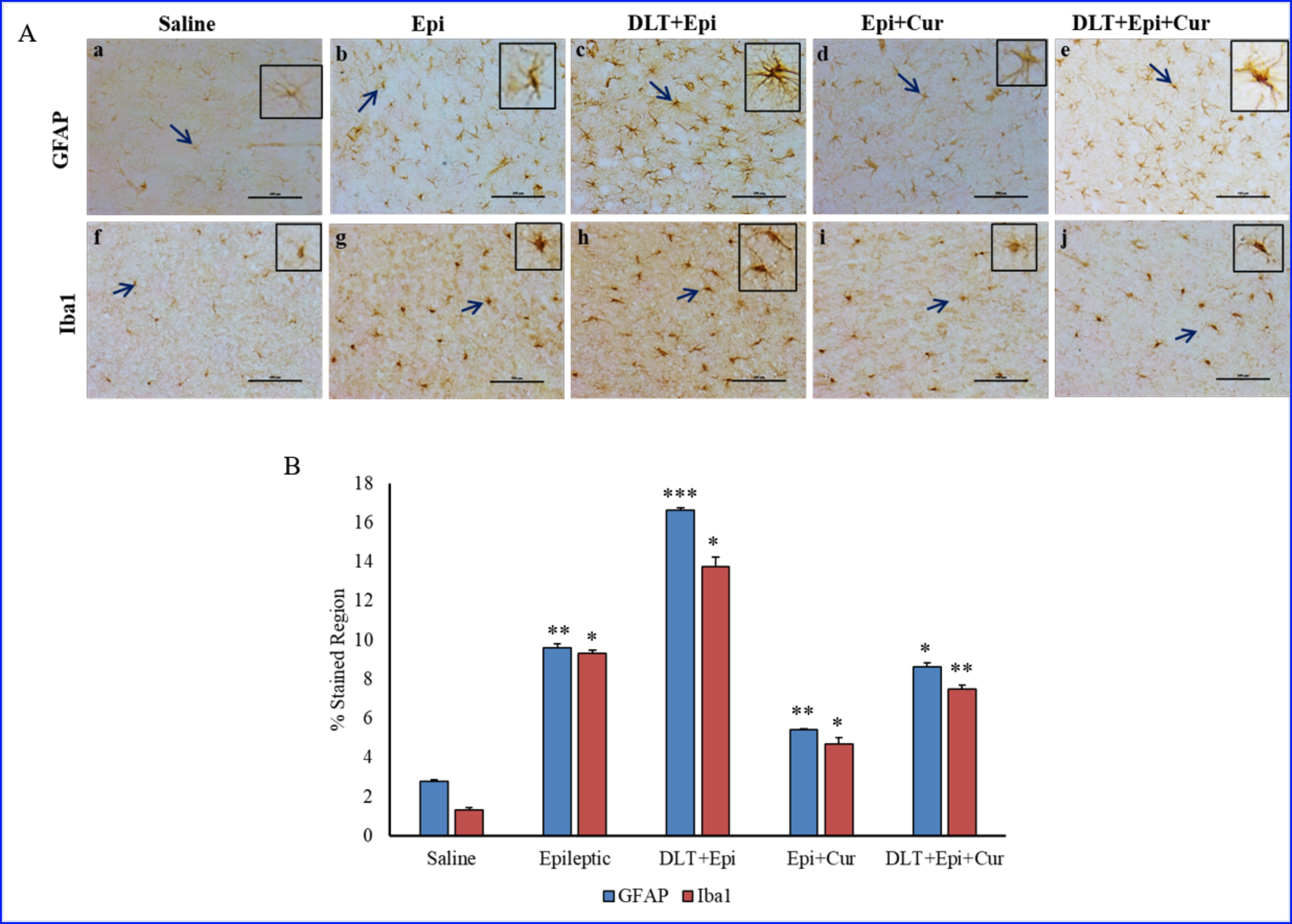
Immunohistology of mouse anti-GFAP and mouse anti-Iba1 in the cortex of saline, Epi and DLT+Epi, Epi+Cur and DLT+Epi+Cur treated groups at 30-day post FeCl_3_ injection (a). Quantification of mean intensity of Nav1.1 and Nav1.6 (b). Insets showing the magnified images of double-labeled Nav1.1 and Nav1.6. Nuclei were stained with DAPI (blue). The scale bar = 100μm. Values are presented as the mean ± SEM (n = 6).

### 3.6. Quantification of GFAP and Iba1

GFAP and Iba1 are markers of glial activation. While GFAP is an astrocytic activation marker, Iba1 is an inflammatory marker present in microglial cells. Here in the current study, we observed and quantified the expression of both GFAP and Iba1 in the cortex region of brain. Compared to saline-injected group, the levels of GFAP and Iba1 were significantly raised in epileptic group (*p* < 0.01, 0.05). Additional exposure of DLT with epilepsy in DLT+Epi group depicted a further significant increase in GFAP and Iba1 (*p* < 0.001, 0.05). However, curcumin treatment significantly lowered both protein levels in both epileptic (*p* < 0.01, 0.05) as well as in DLT+Epi+Cur groups (*p* < 0.05, 0.01). Both markers GFAP (*p* < 0.001) and Iba (*p* < 0.01) showed a significant change between all groups when compared with epileptic group and DLT+Epi group consecutively. However, the DLT+Epi+Cur group shows no significant change Compared to the epileptic group. Similarly, DLT+Epi+Cur shows a significantly higher GFAP and iba1 level when compared to Epi+Cur group (*p* < 0.01), showing curcumin was unable to alleviate the DLT-mediated further rise in levels of both proteins (Fig. 7a and b).

## 4. Discussion

The current study revealed the effects and susceptibility of gestational DLT exposure in adult post-traumatic epileptogenesis. It also finds the decreased antiepileptic effects of curcumin in gestationally DLT-exposed adult epileptic individuals. In particular, we found that FeCl_3_-induced epileptic rats which were exposed to DLT during their embryonic stage through exposing their pregnant mothers showed an increased seizure with raised ECoG firing, MUA count, change in behavioral pattern, decreased PSD95 and SYP, increased neuronal spine density, dendritic complexity, elevated sodium channel subunits, GFAP, and Iba1 expression. We also demonstrated that dietary curcumin treatment alleviated PTE-associated effects. Though the effects of curcumin showed its antiepileptic effects in DLT-exposed rats, the mitigating characteristic diminished.

After a brain trauma, the development of recurrent seizures is a complex process that involves several cellular and molecular changes. These changes may include oxidative damage, changes in neuronal morphology, alterations in axonal and dendritic plasticity, changes in channels and gliosis that subsequently can contribute to the development of seizures [8,11,26]. In the current study, we induced epilepsy by cortical FeCl_3_ injection in rats that were gestationally DLT exposed. After the administration of an iron injection, a singular focal area of the brain becomes the source of epileptic discharges. The discharge from the focus proceeds to spread throughout the entire cerebral cortex and subcortical structures, impacting a broad range of neurological functions [27]. Hence in the current study, the cortical region of the brain post-iron injection was emphasized. Since, the neurotoxic effects of DLT exposure in brain development are also known now [31]. Besides, DLT exposure to neurons is also linked with hyperexcitation to neurons [32]. Hence, our study demonstrated the increase in seizures in gestationally DLT-exposed rats indicating the increased susceptibility to epilepsy after TBI. We also assessed the efficacy of curcumin against DLT-exposed epilepsy-induced rats. [21,33–35]. Thus our study demonstrated reduced epileptic seizures and strong antiepileptic effects. Our findings of curcumin’s antiepileptic effects are in line with the number of previous studies [20,21,36]. However, due to further increased neuronal firing by hyperexcited neurons in DLT-exposed rats, the antiepileptic effect of curcumin is comparably weaker than those without DLT exposure. This clearly indicates that strong DLT neurotoxic comorbid effects are difficult to alleviate with curcumin.

Remarkably, the behavioral alterations due to epilepsy, including anxiety-like behavior and reduced sociability, are well known [9,37,38]. The data from the current study corresponds to the previous reports. The open field maze clearly indicated stress and anxiety through increased traveling time and hyper-exploration in PTE-induced animals. Additionally, current investigation indicates that gestational DLT exposure in iron-induced epileptic rats further reinforced such behavioral alterations due to epilepsy. These behavioral alterations are a consequence of hyperexcitation, inflammation, and morphological changes in neurons [11,20]. Electrophysiology also supported these behavioral changes. Although curcumin is well known to mitigate these epilepsy-associated behavioral alterations, the effects were lesser in DLT-exposed rats when compared to epileptic rats, which were not DLT exposed. This hints at the prenatal neurotoxic effects, which affected epilepsy-associated behavior further, and animals are more prone to higher severity of epilepsy [39,40].

Excitatory synaptic neurons rely on dendritic arbors and spines, which are vital morphological features. Alterations to these structures have been linked to a range of neurological conditions [26]. Specifically, modifications to dendritic arbors and spines have been shown to impact epileptogenesis and the onset of seizures [26,41]. It has been observed that the degeneration of dendrites is a common trait in epileptic neurons [42]. In the future, the stabilization of dendritic structure could serve as a potential therapeutic approach for epilepsy. Though it is established that dendritic arborization and decreased spine density during epilepsy, the mechanism of PTE is yet to be fully understood. In present study is also suggesting similar morphological changes in neurons. Additionally, gestational DLT exposure also strengthened the pieces of evidence we found on neurotoxic effects on susceptibility of PTE.

The mechanism of epileptogenesis consists of multiple factors. The electrophysiological and morphological changes in neurons are a consequence of various cellular events during epileptogenesis. These changes consist of alteration in expression of proteins involved in synapsis, neuronal plasticity and neuronal structural integrity [43]. Notably, neuronal injury and neurological disorders such as epilepsy lead to a decline in synapses which may further lead to neurodegeneration [44,45]. In our study, the reduction in level of PSD95 along with SYP suggested the reinforcement of neurotoxic effects due to gestation DLT exposure causing neuronal damage and a further decrease in synapses. PSD95 and SYP are known markers for synapsis. PSD95 shows the presence of postsynaptic neurons, while SYP provides information about presynaptic neurons. In addition, curcumin is known to reverse these neurodegeneration and synapse declines. Albeit, it shows a similar response to the changes, due to DLT exposure-associated neurotoxicity, curcumin was unable to fully recover the epileptic effects.

The sodium channels NaV1.1 and NaV1.6 are the major voltage-gated channels in mammalian brain, encoded by the genes SCN1A and SCN8A, respectively, involved in the regulation, initiation and propagation of action potentials. These channels are comprised of one alpha and one or two beta subunits [46]. NaV1.6 is known to be present in nods of Ranvier, dendrites and synapses while NaV 1.1 is present in sodium channels of motor neurons [47]. NaV1.1 and NaV 1.6 have been reported to be involved and upregulated in epilepsy [8,36]. Additionally, curcumin also mitigated these epilepsy associated changes in sodium channel subunits [36]. In present study, our results are in line with the present literature. Results also indicate the role of sodium channel subunits in decrease in synapses. Since both Nav1.1 and 1.6 are involved in synaptic transmission [48]. The decrease in pre and postsynaptic neurons could be a leading cause of dysregulation of NaVs subunits. Additionally, the developmental neurotoxic effects of DLT in DLT-exposed rats indicate further enhanced effects similar to previous results. Ameliorative effects of curcumin on epilepsy also function against DLT-mediated neurotoxicity however, the effects are smaller.

Neuroinflammation and Glial activation are characteristics of epilepsy and PTE [26]. GFAP is a known marker of astrocytic activation which occur during neuroinflammation [49]. Similarly, Iba1 is present in microglia and is an indicator of neuroinflammation [50]. Astrogliosis is associated with an increase in glutamate levels in hippocampal sclerosis, leading to heightened neuronal excitability and the occurrence of seizures [51]. Additionally, the reactivation of astrocytes leads to a rapid reduction in the synapses of GABAergic neurons [51]. This suggests that neuronal excitability and inflammation influence the mechanism of reactive glial cells contributing to epileptogenesis. Previous studies have provided substantial evidence highlighting the significance of astroglial activation in epilepsy [26]. In the present study, a significantly higher number of cells positive for GFAP, a marker of astrocyte activation, were observed in the cortex of epileptic rats. The developmental DLT exposure further aggravated these neuroinflammatory responses. Curcumin showed its neuroprotective effects on diminishing neuroinflammation. However, DLT exposure dwindled the curcumin’s antiepileptic effects on rats.

In conclusion, TBI causes an increased stress-mediated response in neurons. Consequently, pathogenesis events of epilepsy appear to be linked with the upregulation of sodium channel subunits causing hyperexcitation. This neuronal hyperexcitation due to upregulation of NaVs may further lead to neuroinflammation, synaptic decline, decreased dendritic spine and increased neurodegeneration. Such intra and intercellular event cause disrupted behavioral responses. Though DLT is unable to cross BBB, it can enter the developing brain of an embryo. DLT may hyperexcited neurons further. Hence, it is possible that gestational DLT exposure raised the susceptibility to epilepsy, aggravated epilepsy-mediated alterations and diminished effects of curcumin in the PTE.

## 7. Conflict of Interest

The authors declare that the research was conducted in the absence of any commercial or financial relationships that could be construed as a potential conflict of interest. The author declares that there is no conflict of interest.

## 8. Author Contributions

PK and KK contributed equally in study concept, study design, interpretation of results and manuscript preparation. Author DS contributed in assessment of results and manuscript preparation. All authors commented on previous versions of the manuscript. All authors read and approved the final manuscript.

## 9. Funding

Department of Biotechnology, Gov. of India provided funding for this work.

## 10. Acknowledgements

Authors are thankful to Department of Biotechnology, Gov. of India for providing funding and Jawaharlal Nehru University of providing infrastructure for this work.

## Notes

### Competing Interest Statement

The authors have declared no competing interest.

## 5. References

1. Lu Q, Sun Y, Ares I, Anadón A, Martínez MMA, Martínez-Larrañaga MR, et al. Deltamethrin toxicity: A review of oxidative stress and metabolism. Environ Res [Internet]. 2019 [cited 2023 Mar 13];170:260–81. Available from: 10.1016/j.envres.2018.12.045

2. Chrustek A, Hołyńska-Iwan I, Dziembowska I, Bogusiewicz J, Wróblewski M, Cwynar A, et al. Current research on the safety of pyrethroids used as insecticides. Med. 2018;54:1–15.

3. Sharma P, Singh R, Jan M. Dose-dependent effect of deltamethrin in testis, liver, and Kidney of wistar rats. Toxicol Int. 2014;21:131–9.

4. Lei L, Zhu B, Qiao K, Zhou Y, Chen X, Men J, et al. New evidence for neurobehavioral toxicity of deltamethrin at environmentally relevant levels in zebrafish. Sci Total Environ [Internet]. 2022;822:153623. Available from: 10.1016/j.scitotenv.2022.153623

5. Patro N, Shrivastava M, Tripathi S, Patro IK. S100β upregulation: A possible mechanism of deltamethrin toxicity and motor coordination deficits. Neurotoxicol Teratol. 2009;31:169– 76.

6. Kumar K, Patro N, Patro I. Impaired structural and functional development of cerebellum following gestational exposure of deltamethrin in rats: Role of reelin. Cell Mol Neurobiol [Internet]. 2013 [cited 2023 Jul 25];33:731–46. Available from: https://link.springer.com/article/10.1007/s10571-013-9942-7

7. Park JT, Delozier SJ, Chugani HT. Epilepsy due to mild TBI in children: An experience at a tertiary referral center. J Clin Med. 2021;10:1–12.

8. Prakash C, Mishra M, Kumar P, Kumar V, Sharma D. Response of Voltage-Gated Sodium and Calcium Channels Subtypes on Dehydroepiandrosterone Treatment in Iron-Induced Epilepsy. Cell Mol Neurobiol [Internet]. 2021 [cited 2021 Dec 13];41:279–92. Available from: https://link.springer.com/article/10.1007/s10571-020-00851-0

9. Castelhano ASS, Cassane G dos ST, Scorza FA, Cysneiros RM. Altered anxiety-related and abnormal social behaviors in rats exposed to early life seizures. Front Behav Neurosci. 2013;7:1–8.

10. Srivastava NK, Mukherjee S, Sharma R, Das J, Sharma R, Kumar V, et al. Altered lipid metabolism in post-traumatic epileptic rat model: one proposed pathway. Mol Biol Rep [Internet]. 2019;46:1757–73. Available from: 10.1007/s11033-019-04626-9

11. Mishra M, Singh R, Sharma D. Antiepileptic action of exogenous dehydroepiandrosterone in iron-induced epilepsy in rat brain. Epilepsy Behav [Internet]. 2010;19:264–71. Available from: 10.1016/j.yebeh.2010.06.048

12. Nelson KM, Dahlin JL, Bisson J, Graham J, Pauli GF, Walters MA. The Essential Medicinal Chemistry of Curcumin. J Med Chem [Internet]. 2017;60:1620–37. Available from: 10.1021/acs.jmedchem.6b00975

13. Aggarwal BB, Harikumar KB. Potential Therapeutic Effects of Curcumin, the Anti-inflammatory Agent, Against Neurodegenerative, Cardiovascular, Pulmonary, Metabolic, Autoimmune and Neoplastic Diseases. Int J Biochem Cell Biol [Internet]. 2009;41:40–59. Available from: http://linkinghub.elsevier.com/retrieve/pii/S1357272508002550

14. Kulkarni S, Dhir A. An overview of curcumin in neurological disorders. Indian J Pharm Sci [Internet]. 2010;72:149. Available from: http://www.ijpsonline.com/text.asp?2010/72/2/149/65012

15. Wang P, Su C, Li R, Wang H, Ren Y, Sun H, et al. Mechanisms and effects of curcumin on spatial learning and memory improvement in APPswe/PS1dE9 mice. J Neurosci Res [Internet]. 2014 [cited 2018 Jun 6];92:218–31. Available from: http://doi.wiley.com/10.1002/jnr.23322

16. Benameur T, Soleti R, Panaro MA, La Torre ME, Monda V, Messina G, et al. Curcumin as prospective anti-aging natural compound: Focus on brain. Molecules. 2021;26:1–17.

17. Huang HC, Chang P, Dai XL, Jiang ZF. Protective effects of curcumin on amyloid-b-induced neuronal oxidative damage. Neurochem Res [Internet]. 2012 [cited 2023 Jul 13];37:1584–97. Available from: https://link.springer.com/article/10.1007/s11064-012-0754-9

18. Tripanichkul W, Jaroensuppaperch EO. Curcumin protects nigrostriatal dopaminergic neurons and reduces glial activation in 6-hydroxydopamine hemiparkinsonian mice model. Int J Neurosci [Internet]. 2012 [cited 2023 Jul 13];122:263–70. Available from: https://pubmed.ncbi.nlm.nih.gov/22176529/

19. Dhir A. Curcumin in epilepsy disorders. Phyther Res [Internet]. 2018 [cited 2023 Jul 13];32:1865–75. Available from: https://onlinelibrary.wiley.com/doi/full/10.1002/ptr.6125

20. Jyoti A, Sethi P, Sharma D. Curcumin protects against electrobehavioral progression of seizures in the iron-induced experimental model of epileptogenesis. Epilepsy Behav. 2009;14:300–8.

21. Kumar P, Sharma D. Ameliorative effect of curcumin on altered expression of CACNA1A and GABRD in the pathogenesis of FeCl3-induced epilepsy. Mol Biol Rep [Internet]. 2020 [cited 2023 Mar 14];47:5699–710. Available from: 10.1007/s11033-020-05538-9

22. Kumar P, Singh R, Sharma D. ALTERED EXPRESSION OF MIR-214, MIR-3120 AND PTEN IN IRON-INDUCED EXPERIMENTAL EPILEPSY MODEL OF POST-TRAUMATIC EPILEPSY AND THE EFFECT OF CURCUMIN. Int J Adv Res [Internet]. 2016 [cited 2018 Jun 11];4:1352–61. Available from: http://www.journalijar.com/article/14035/altered-expression-of-mir-214,-mir-3120-and-pten-in-iron-induced-experimental-epilepsy-model-of-post-traumatic-epilepsy-and-the-effect-of-curcumin./

23. Kandratavicius L, Alves Balista P, Lopes-Aguiar C, Ruggiero RN, Umeoka EH, Garcia-Cairasco N, et al. Animal models of epilepsy: Use and limitations [Internet]. Neuropsychiatr. Dis. Treat. Dove Medical Press Ltd.; 2014 [cited 2020 Aug 26]. p. 1693–705. Available from: /pmc/articles/PMC4164293/?report=abstract

24. Wang Y, Wei P, Yan F, Luo Y, Zhao G. Animal Models of Epilepsy: A Phenotype-oriented Review [Internet]. Aging Dis. International Society on Aging and Disease; 2022 [cited 2023 Jul 13]. p. 215–31. Available from: https://www.aginganddisease.org/EN/10.14336/AD.2021.0723

25. Mosini AC, Calió ML, Foresti ML, Valeriano RPS, Garzon E, Mello LE. Modeling of post-traumatic epilepsy and experimental research aimed at its prevention. Brazilian J Med Biol Res [Internet]. 2020 [cited 2023 Jul 13];54:1–12. Available from: /pmc/articles/PMC7747873/

26. Prakash C, Rabidas SS, Tyagi J, Sharma D. Dehydroepiandrosterone Attenuates Astroglial Activation, Neuronal Loss and Dendritic Degeneration in Iron-Induced Post-Traumatic Epilepsy. Brain Sci [Internet]. 2023 [cited 2023 Jul 13];13:563. Available from: https://www.mdpi.com/2076-3425/13/4/563/htm

27. Mishra M, Singh R, Mukherjee S, Sharma D. Dehydroepiandrosterone’s antiepileptic action in FeCl3-induced epileptogenesis involves upregulation of glutamate transporters. Epilepsy Res [Internet]. 2013 [cited 2023 Jul 13];106:83–91. Available from: 10.1016/j.eplepsyres.2013.06.008

28. Kaidanovich-Beilin O, Lipina T, Vukobradovic I, Roder J, Woodgett JR. Assessment of Social Interaction Behaviors. J Vis Exp [Internet]. 2011 [cited 2023 Jul 13];2473. Available from: /pmc/articles/PMC3197404/

29. Lee E-H, Park J-Y, Lee Y, Han P-L. Sociability and Social Novelty Preference Tests Using a U-shaped Two-choice Field. Bio-protocol [Internet]. 2018 [cited 2023 Jul 13];8. Available from: /pmc/articles/PMC8275281/

30. Patro N, Kumar K, Patro I. Quick Golgi method: Modified for high clarity and better neuronal anatomy. Indian J Exp Biol. 2013;51:685–93.

31. Ding R, Cao Z, Wang Y, Gao X, Luo H, Zhang C, et al. The implication of p66shc in oxidative stress induced by deltamethrin. Chem. Biol. Interact. Elsevier; 2017.

32. Harrill JA, Li Z, Wright FA, Radio NM, Mundy WR, Tornero-Velez R, et al. Transcriptional response of rat frontal cortex following acute In Vivo exposure to the pyrethroid insecticides permethrin and deltamethrin. BMC Genomics [Internet]. 2008 [cited 2023 Jul 13];9:1–23. Available from: https://bmcgenomics.biomedcentral.com/articles/10.1186/1471-2164-9-546

33. Kong Y, Li M, Guo G, Yu L, Sun L, Yin Z, et al. Effects of dietary curcumin inhibit deltamethrin-induced oxidative stress, inflammation and cell apoptosis in Channa argus via Nrf2 and NF-κB signaling pathways. Aquaculture. 2021;540:736744.

34. Sharma P, Singh R. Protective role of curcumin in deltamethrin induced system toxicity in Wistar rats. Planta Med [Internet]. 2013 [cited 2023 Jul 13];79:PB43. Available from: http://www.thieme-connect.de/DOI/DOI?10.1055/s-0033-1351988

35. Zaki SM, Algaleel WAA, Imam RA, Soliman GF, Ghoneim FM. Nano-curcumin versus curcumin in amelioration of deltamethrin-induced hippocampal damage. Histochem Cell Biol [Internet]. 2020 [cited 2023 Jul 13];154:157–75. Available from: https://link.springer.com/article/10.1007/s00418-020-01871-z

36. Kumar V, Prakash C, Singh R, Sharma D. Curcumin’s antiepileptic effect, and alterations in Nav1.1 and Nav1.6 expression in iron-induced epilepsy. Epilepsy Res. 2019;150:7–16.

37. Kleen JK, Scott RC, Lenck-Santini P-P, Holmes GL. Cognitive and Behavioral Co-Morbidities of Epilepsy. Jasper’s Basic Mech Epilepsies [Internet]. 2012 [cited 2023 Jul 14];915–29. Available from: https://www.ncbi.nlm.nih.gov/books/NBK98139/

38. Vazquez B, Devinsky O. Epilepsy and anxiety. Epilepsy Behav [Internet]. 2003 [cited 2023 Jul 13];4:20–5. Available from: http://www.epilepsybehavior.com/article/S1525505003002841/fulltext

39. Lazarini CA, Florio JC, Lemonica IP, Bernardi MM. Effects of prenatal exposure to deltamethrin on forced swimming behavior, motor activity, and striatal dopamine levels in male and female rats. Neurotoxicol Teratol. 2001;23:665–73.

40. Husain R, Husain R, Vaqar M, Adhami PK, Seth; K, Behavioral N, et al. BEHAVIORAL, NEUROCHEMICAL, AND NEUROMORPHOLOGICAL EFFECTS OF DELTAMETHRIN IN ADULT RATS. 101080/009841096161212 [Internet]. 2010 [cited 2023 Jul 14];48:515–6. Available from: https://www.tandfonline.com/doi/abs/10.1080/009841096161212

41. Teskey GC, Monfils MH, Silasi G, Kolb B. Neocortical kindling is associated with opposing alterations in dendritic morphology in neocortical layer V and striatum from neocortical layer III. Synapse [Internet]. 2006 [cited 2023 Jul 14];59:1–9. Available from: https://pubmed.ncbi.nlm.nih.gov/16235229/

42. Rossini L, de Santis D, Mauceri RR, Tesoriero C, Bentivoglio M, Maderna E, et al. Dendritic pathology, spine loss and synaptic reorganization in human cortex from epilepsy patients. Brain [Internet]. 2021 [cited 2023 Jul 15];144:251–65. Available from: https://pubmed.ncbi.nlm.nih.gov/33221837/

43. Bernard C. Alterations in synaptic function in epilepsy. Epilepsia [Internet]. 2012 [cited 2023 Jul 15];51:42. Available from: https://www.ncbi.nlm.nih.gov/books/NBK98161/

44. do Canto AM, Donatti A, Geraldis JC, Godoi AB, da Rosa DC, Lopes-Cendes I. Neuroproteomics in Epilepsy: What Do We Know so Far? Front Mol Neurosci. 2021;13:604158.

45. Ye Q, Zeng C, Luo C, Wu Y. Ferrostatin-1 mitigates cognitive impairment of epileptic rats by inhibiting P38 MAPK activation. Epilepsy Behav. 2020;103:106670.

46. Catterall WA. From ionic currents to molecular mechanisms: the structure and function of voltage-gated sodium channels. Neuron [Internet]. 2000 [cited 2023 Jul 15];26:13–25. Available from: https://pubmed.ncbi.nlm.nih.gov/10798388/

47. Caldwell JH, Schaller KL, Lasher RS, Peles E, Levinson SR. Sodium channel Nav1.6 is localized at nodes of Ranvier, dendrites, and synapses. Proc. Natl. Acad. Sci. U. S. A. 2000. p. 5616–20.

48. Vysokov N, McMahon SB, Raouf R. The role of NaV channels in synaptic transmission after axotomy in a microfluidic culture platform. Sci Rep [Internet]. 2019 [cited 2023 Jul 15];9:12915. Available from: /pmc/articles/PMC6733904/

49. Shimada T, Takemiya T, Sugiura H, Yamagata K. Role of inflammatory mediators in the pathogenesis of epilepsy. Mediators Inflamm. 2014;2014.

50. Sharma S, Tiarks G, Haight J, Bassuk AG. Neuropathophysiological Mechanisms and Treatment Strategies for Post-traumatic Epilepsy. Front Mol Neurosci. 2021;14.

51. Ortinski PI, Dong J, Mungenast A, Yue C, Takano H, Watson DJ, et al. Selective induction of astrocytic gliosis generates deficits in neuronal inhibition. Nat Neurosci [Internet]. 2010 [cited 2023 Jul 15];13:584–91. Available from: https://pubmed.ncbi.nlm.nih.gov/20418874/

